# SARS-CoV-2 proteome microarray for mapping COVID-19 antibody interactions at amino acid resolution

**DOI:** 10.1101/2020.03.26.994756

**Authors:** Hongye Wang, Xin Hou, Xian Wu, Te Liang, Xiaomei Zhang, Dan Wang, Fei Teng, Jiayu Dai, Hu Duan, Shubin Guo, Yongzhe Li, Xiaobo Yu

## Abstract

COVID-19 has quickly become a worldwide pandemic, which has significantly impacted the economy, education, and social interactions. Understanding the humoral antibody response to SARS-CoV-2 proteins may help identify biomarkers that can be used to detect and treat COVID-19 infection. However, no immuno-proteomics platform exists that can perform such proteome-wide analysis. To address this need, we created a SARS-CoV-2 proteome microarray to analyze antibody interactions at amino acid resolution by spotting peptides 15 amino acids long with 5-amino acid offsets representing full-length SARS-CoV-2 proteins. Moreover, the array processing time is short (1.5 hours), the dynamic range is ~2 orders of magnitude, and the lowest limit of detection is 94 pg/mL. Here, the SARS-CoV-2 proteome array reveals that antibodies commercially available for SARS-CoV-1 proteins can also target SARS-CoV-2 proteins. These readily available reagents could be used immediately in COVID-19 research. Second, IgM and IgG immunogenic epitopes of SARS-CoV-2 proteins were profiled in the serum of ten COVID-19 patients. Such epitope biomarkers provide insight into the immune response to COVID-19 and are potential targets for COVID-19 diagnosis and vaccine development. Finally, serological antibodies that may neutralize viral entry into host cells via the ACE2 receptor were identified. Further investigation into whether these antibodies can inhibit the propagation of SARS-CoV-2 is warranted. Antibody and epitope profiling in response to COVID-19 is possible with our peptide-based SARS-COV-2 proteome microarray. The data gleaned from the array could provide invaluable information to the scientific community to understand, detect, and treat COVID-19.

---

The first case of infection caused by the novel coronavirus (SARS-CoV-2) was reported in Wuhan City, China, in December 2019. SARS-CoV-2 has since proven to be highly infectious, with the median incubation period of 4 days ^1–3^. Infection of SARS-CoV-2, called COVID-19, results in a range of symptoms, ranging from a mild cough to pneumonia. It is estimated that 17.9% of patients might be asymptomatic^4^, which may lead to two or even three transmissions per infected individual ^3,5,6^. Particular subsets of the population are extremely vulnerable to COVID-19, including the elderly, those with underlying conditions, and immunocompromised individuals. For example, 80% of the deaths attributed to COVID-19 occur among adults ≥ 65 years old. On the evening of January 30, 2020, the World Health Organization listed the novel coronavirus outbreak as a public health emergency of international concern ^7^. As of March 25, the novel coronavirus had spread worldwide^8^, with 417,966 confirmed cases and 18,615 deaths in 169 countries. The high transmission rates of SARS-CoV-2, limited diagnostic tests, and no anti-viral treatment options pose huge challenges for the control and treatment of SARS-CoV-2 infected patients ^9,10^.

SARS-CoV-2 is 82% similar to the original SARS virus attributed to the outbreak in 2003^11^. Generally, a mature SARS-CoV-2 virus has a polyprotein (the open reading frame 1a and 1b, Orf1ab), four structural proteins (envelope, E; membrane, M; nucleocapsid, N; spike, S) and five accessary proteins (Orf3a, Orf6, Orf7a, Orf8, Orf10)^12^. The Orf1ab is involved in viral RNA replication and transcription^12^. The E and M proteins are important in the viral assembly of a coronavirus, and the N protein is necessary for viral RNA synthesis^13^. The S protein is on the surface of the viral particle, enabling the infection of host cells by binding to the host cell receptor, ACE2, via the S-protein’s receptor binding domain (RBD) within the S-protein’s subunit 1. The RBD of SARS-CoV-2 is very different from the S protein’s RBD of SARS-CoV-1; in fact, they only share 73.6% homology^14^. The SARS-CoV-2 RBD can bind faster to the ACE2 receptor than the SARS-CoV-1 RBD, thus resulting in high transmission efficiency of the virus^14^. The accessory proteins may have functions in signaling inhibition, apoptosis induction and cell cycle arrest^12^.

B cells defend the body against viruses, such as the SARS-CoV-2 virus, by producing antibodies, which bind to viral particles to mark them for destruction by other cells in the immune system ^15,16^. However, there still don’t have an immuno-proteomics platform to perform proteome-wide analysis of humoral antibody response to SARS-CoV-2 proteins yet. Here we describe a SARS-CoV-2 proteome peptide microarray that enables high throughput antibody screening of COVID-19 patients to all SARS-CoV-2 protein sequences at amino acid resolution.

To produce the SARS-CoV-2 proteome microarray (Figure 1a), we first extracted the reference sequences of ten proteins encoded by the SARS-CoV-2 coronavirus genome from the NCBI database (<u>Accession no. MN908947.3</u>). Using these reference sequences, we prepared a peptide library containing 966 peptides representing SARS-CoV-2 proteins, in which each peptide was 15 amino acids long with a 5 amino acid overlap. All peptides were labeled with a C-terminal biotin group and printed onto a 3D-modified microscope slide using biotin-streptavidin chemistry. Full-length SARS-COV-2 N protein, full-length E, and five S truncated proteins were also printed. (Supplementary Table 1).

**Figure 1.**
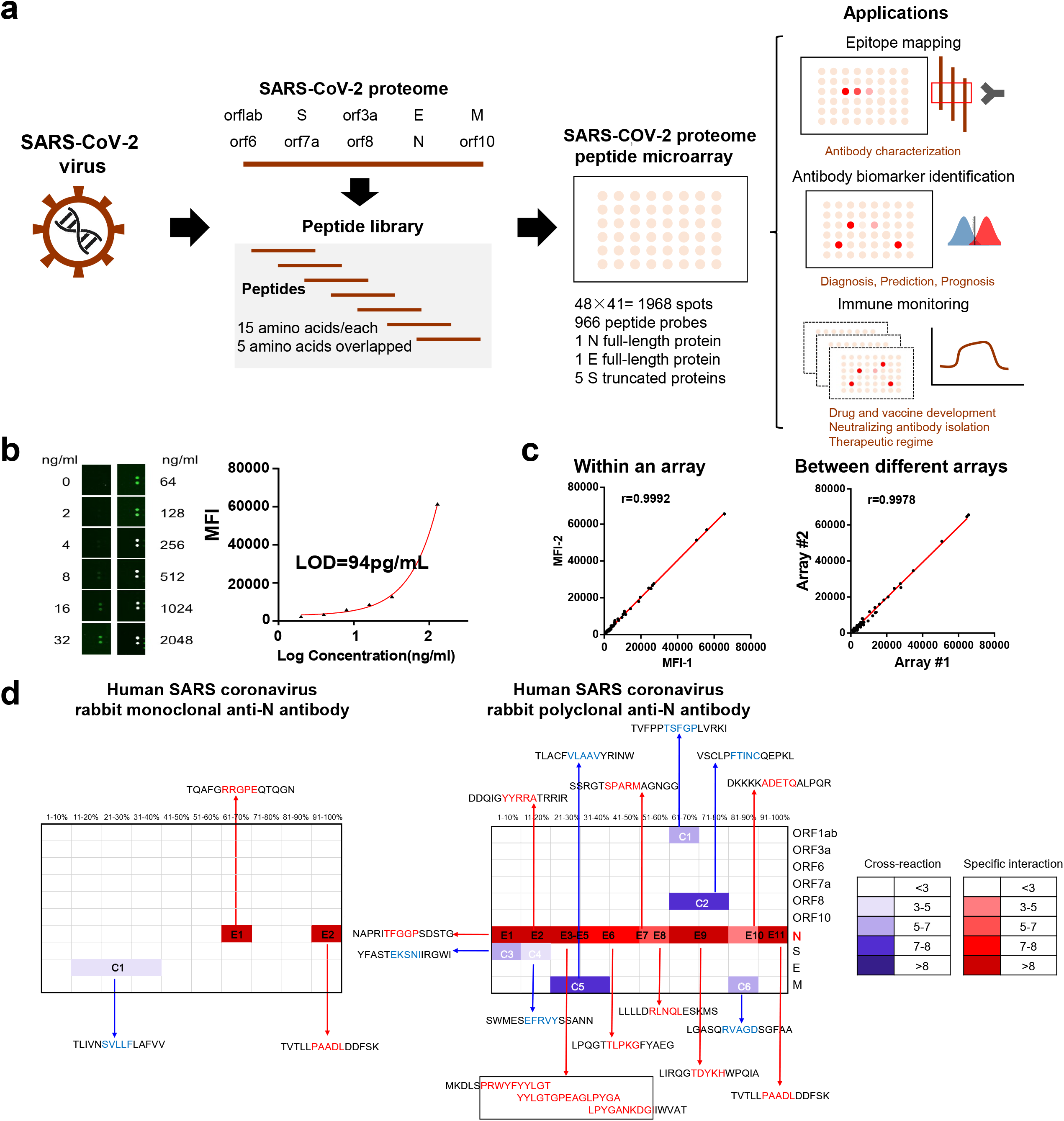
SARS-COV-2 proteome microarray fabrication and application in antibody characterization. (a) The schematic illustration of SARS-COV-2 proteome microarray fabrication and biomedical applications; (b) Dynamic range of serum antibody detection using SARS-COV-2 proteome microarray. The lowest of detection limit (LOD) was calculated using the signal of the buffer control plus two standard deviations. (c) Reproducibility of serum antibody detection using the SARS-COV-2 proteome microarray. (c) Epitope binding of the anti-SARS-CoV-1 N protein antibody using the SARS-COV-2 proteome microarray. The specific antibody binding to the target epitope is selected with a Z-score higher than 3 as a threshold. The false-colored rainbow color from blue to red corresponds to the Z-score from low to high, respectively.

We next determined the optimal lengths of time to block the array, incubate with serum samples, and incubate with the detection antibody using serum spiked with anti-SARS antibodies. Optimal signal-to-noise ratios were obtained with blocking for 1 min, serum incubation for 30 min, and detection antibody incubation for 30 min (Supplemental Figures 1 – 3). Serum screening using the SARS-CoV-2 proteome microarray can be performed in 1.5 hours while keeping a good dynamic range (~2 orders) and sensitivity (94 pg/mL) (Figure 1b). This represents a significant decrease in time compared to the standard ~ 18 hours using protein microarrays ^17^. The r correlation within an array and between different arrays were 0.9992 and 0.9978, respectively, demonstrating the high reproducibility the SARS-COV-2 proteome microarrays (Figure 1c).

Since the SARS-CoV-1 and SARS-CoV-2 genomes are highly similar, we tested rabbit monoclonal and polyclonal anti-SARS-CoV-1 N protein antibodies on the SARS-COV-2 proteome microarray (Figure 1d, Supplementary Figure 4). The monoclonal Ab displayed high specificity to two epitopes (RRGPE and PAADL) on the SARS-COV-2 N protein with a Z-score higher than 3. The functions of these epitopes are currently unknown. Minor cross-reactivity was observed on the epitope (SVLLF) of the E protein. The polyclonal antibody bound to eleven epitopes (E1-E11) on the N protein with crossreactivity to six epitopes on M, S, ORF8, and ORF1ab proteins. The cross-reactive epitopes on M, S, ORF8, and ORF1ab proteins are different than those present in the N-protein (Figure 1d), and it is unclear why the antibodies have crossreactivity to these proteins. The results were validated using full-length N- and S-proteins (Supplementary Figure 5). These results demonstrate that the antibodies prepared previously to SARS-CoV-1 proteins may also detect SARS-COV-2 proteins ^18^. SARS-CoV-1 antibodies could provide a quick alternative to fighting COVID-19 since the generation of antibodies to SARS-COV-2 proteins will take three to six months.

To demonstrate the utility of the SARS-COV-2 proteome microarray in antibody profiling, we screened IgM and IgG antibodies in the serum of ten COVID-19 patients and constructed a landscape of humoral response to the SARS-COV-2 proteome (Figure 2). All IgG and IgA antibody epitopes were identified with a Z-score higher than 3 in at least one of COVID-19 patients (Table 1). Many antibodies targeted peptides from seven SARS-COV-2 proteins (M, N, S, Orf1ab, Orf3a, Orf7a, and Orf8). Notably, four immunodominant epitopes with antibodies in more than 80% COVID-19 patients were present in N (residue 206-210, SPARM), S (residue 816-820, SFIED) and Orf3a (residue 136-140, KNPLL; residue 176-180, SPISE) proteins. However, antibodies to E, Orf6 and Orf10 were not detected.

**Figure 2.**
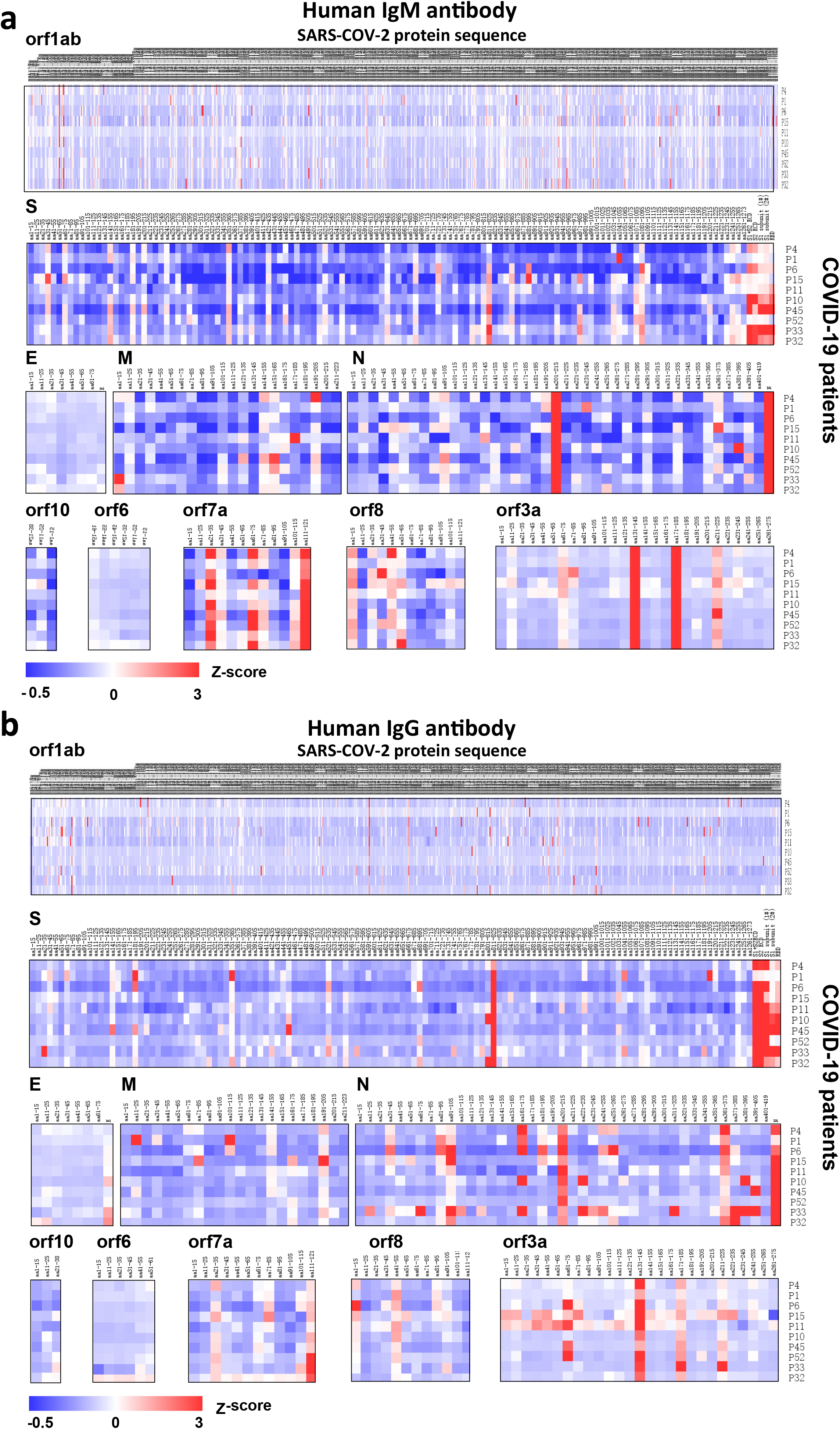
Landscape of humoral antibody response to SARS-COV-2 proteome. (a) and (b) are the distribution of human IgM and IgG antibodies to SARS-COV-2 individual proteins, respectively. The x-axis represents the sequence of amino acids of SARS-COV-2 proteins. The y-axis represents the serum samples from COVID-19 patients. The false-colored rainbow color from blue to red corresponds to the signals of antibody binding from low to high, respectively.

**Table 1.**
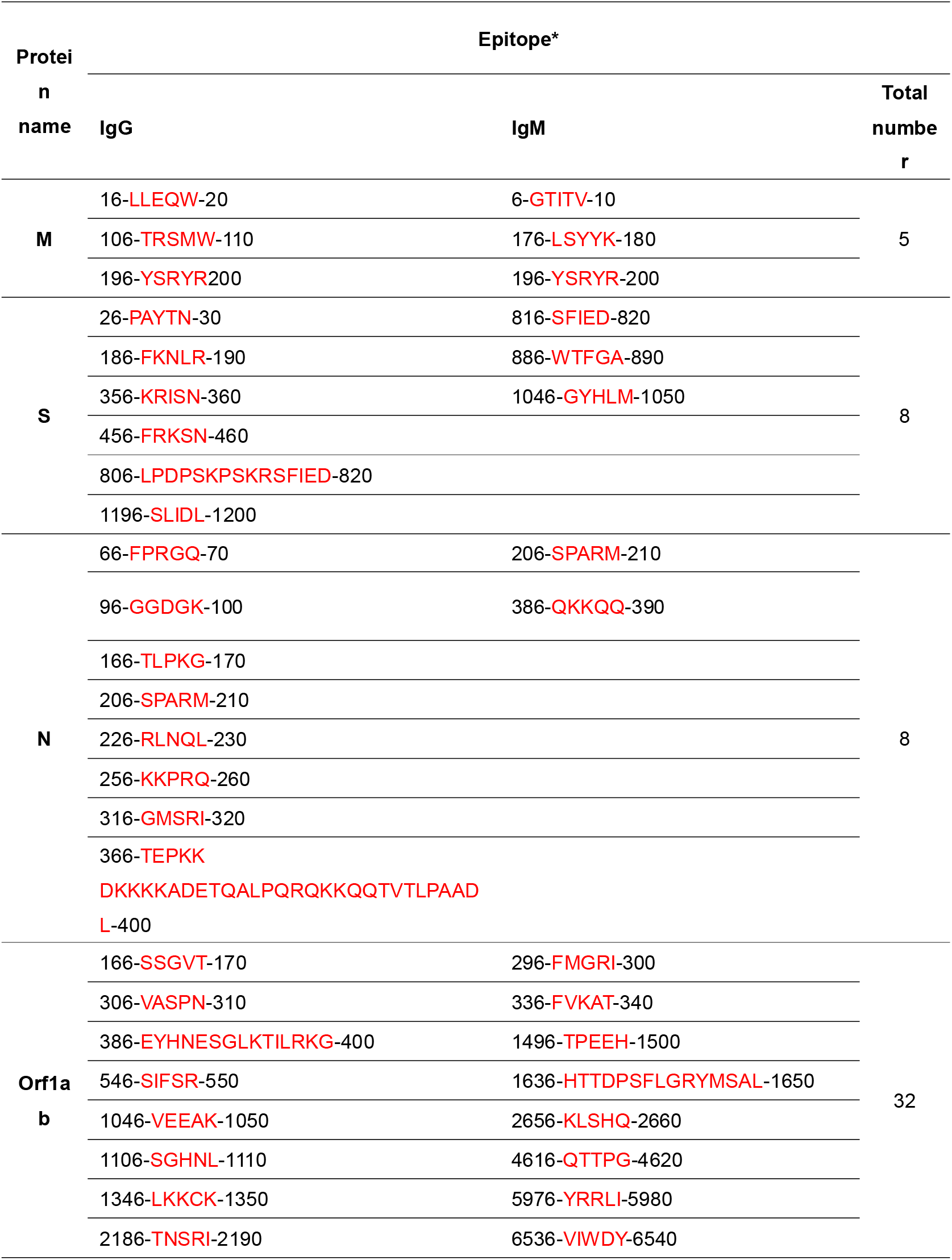
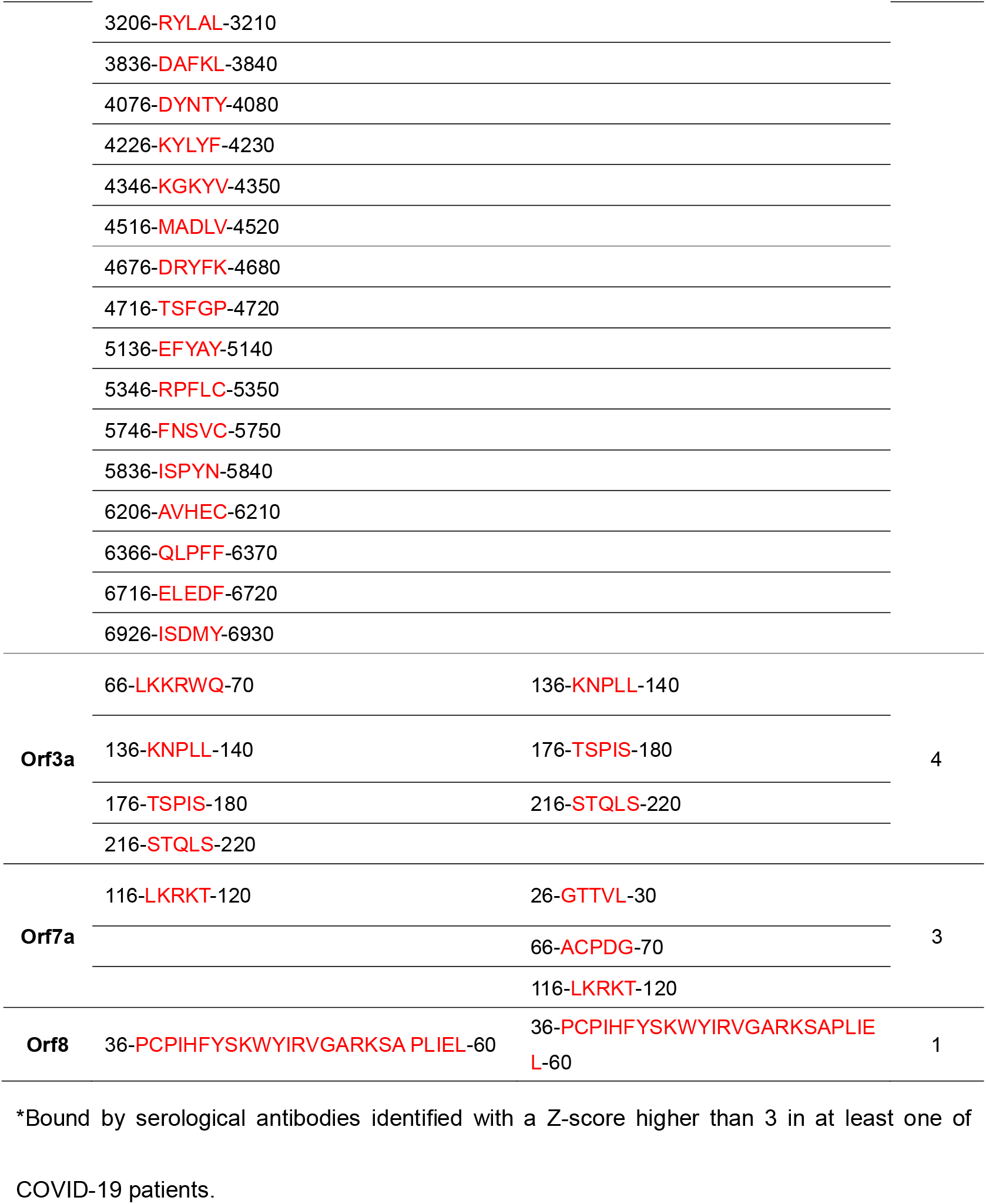
The epitopes identified in the serum of COVID-19 patients using SARS-COV-2 proteome microarrays.

The identification of B-cell and T-cell epitopes for SARS-COV-2 proteins is critical for the development of effective diagnostic tests and vaccine, especially for structural N and S proteins, which had been predicted by bioinformatics^19,20^. For example, there are several RT-PCR tests for detecting COVID-19 that target the S or N protein genome^21^. ELISA and lateral flow devices are also available that measure IgM and IgG levels to N or S proteins^22^. In this work, using overlapping peptides representing the full-length S protein, human IgM and human IgG antibodies were found to target three and six epitopes, respectively (Figure 2, Table 1). Likewise for the N-protein, IgM antibodies targeted two epitopes and IgG antibodies bound to eight epitopes (Figure 2, Table 1). The structural analysis show that all epitope peptides in RNA binding domain of N protein are located at the loop that are easily accessible to the antibodies (Figure 3a). Six epitopes were identified in S protein structure, in which three epitopes are located at the surface and three epitopes are located inside of the protein (Figure 3b). From the predicted B-cell epitopes^19,20^, two epitopes (residue 806-820, LPDPSKPSKRSFIED; residue 456-460, FRKSN)) on S protein, one epitope on N protein (residue 166-170, TLPKG), and one epitope (residue 6-10, GTITV) on M protein were experimentally confirmed in this study. The B-cell epitopes that are identified for SARS-COV-2 proteome using microarray will facilitate the understanding of B-cell immunity, identify biomarkers and vaccine candidates for COVID-19 treatment, which should be investigated in a large clinical cohort in future^20^.

**Figure 3.**
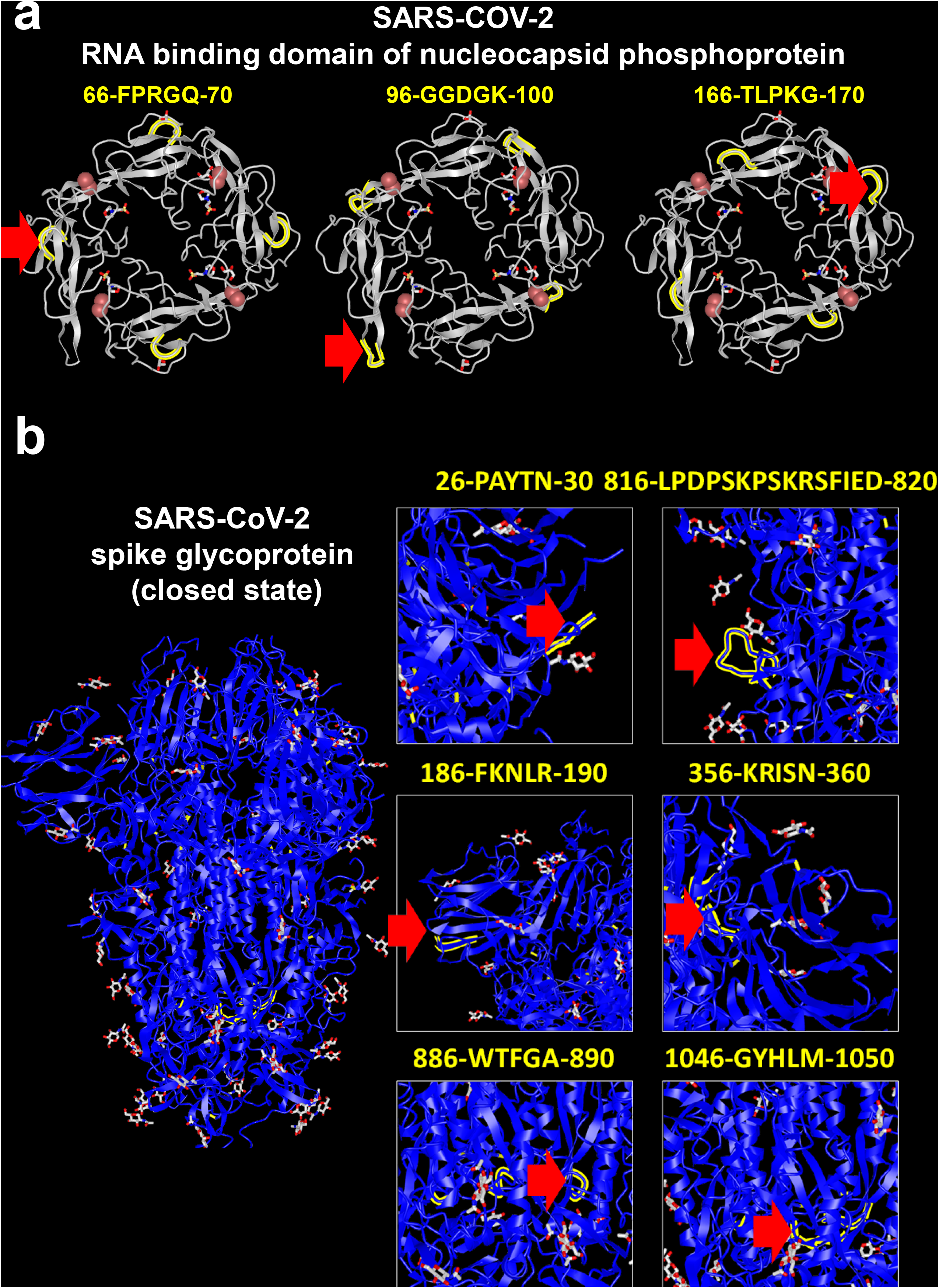
Structural analysis of immunogenic epitopes SARS-COV-2 proteins. (a) and (b) are the structural analysis of nucleocapsid RNA binding domain (PDB ID: 6VYO) and spike trimer protein (PDB ID: 6VXX). The epitope is labeled with yellow and indicated with red arrow.

The SARS-COV-2 S-protein’s RBD (residue 438-498) directly engages the ACE2 receptor and might be an ideal target for developing neutralizing antibodies^23^. However, the identification of neutralizing antibodies to competitively inhibit the binding of SARS-COV-2 virus to the host ACE2 receptor has proved challenging. In this work, we analyzed the immunological response to seven peptide sequences within the RBD. Some IgM antibodies from patient “P52” and IgG antibodies from patients “P10” and “P45” bind to the same epitope (residue 456-460, FRKSN)(Figure 4a and 4b). Structural analysis of the RBD-ACE2 complex shows that the epitope is located in the RBD loop engaged with the ACE2 receptor^24^ (Figure 4c), thus supporting our data. This epitope may serve as an antigen to stimulate neutralizing antibodies to the RBD-ACE2 interaction and increase CD4+/CD8+ T-cell responses^19,25^.

**Figure 4.**
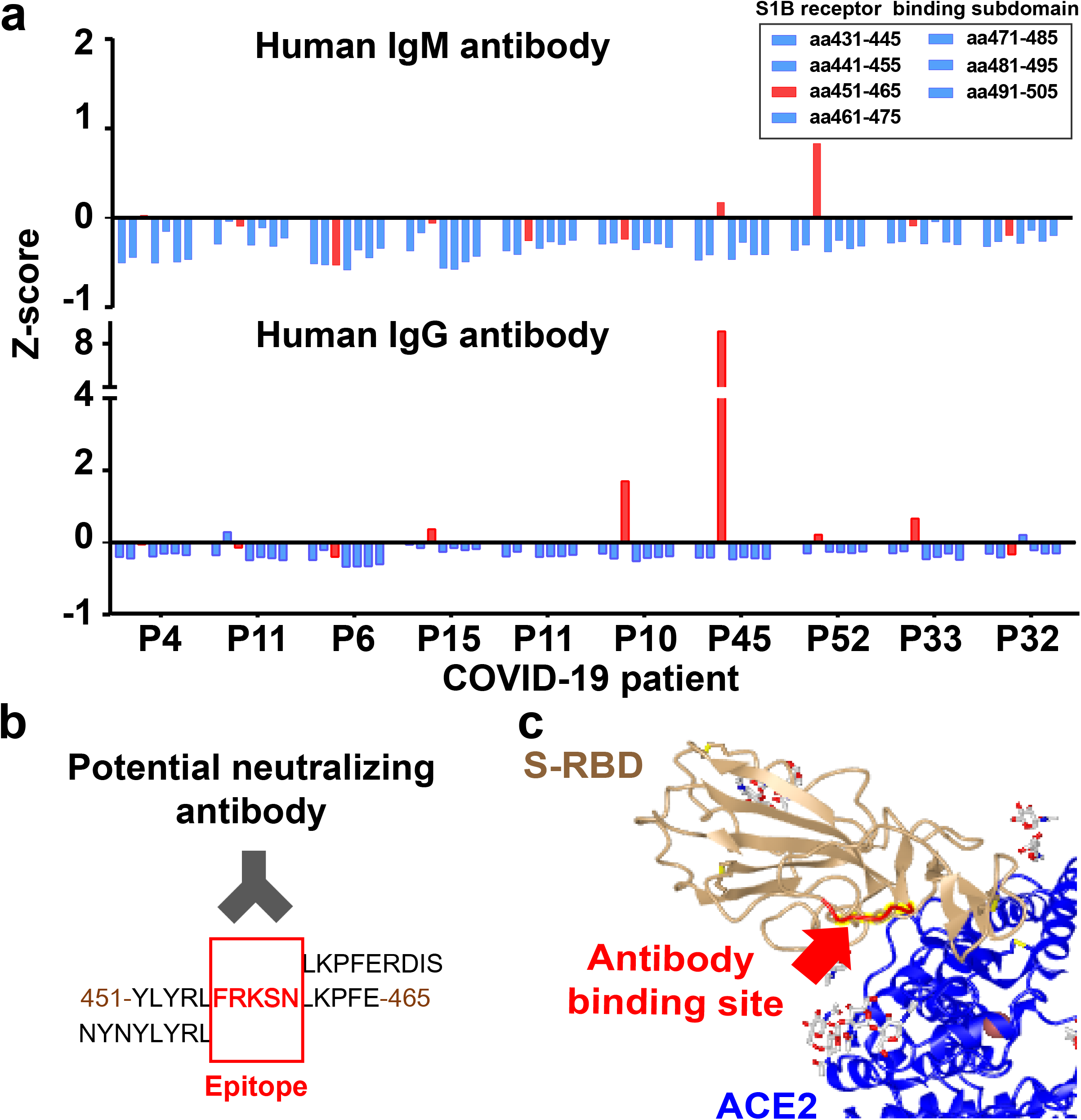
Identification of potential neutralizing antibody targets in the serum of COVID-19 patients using the SARS-COV-2 proteome microarray. (a) Z-score of serum antibody binding to the peptides with the S-proteins RBD (amino acid residues 431-505). (b) Identification of antibody binding epitope (FRKSN) through sequence alignment. (c) Schematic illustration of the epitope on the RBD (FRKSN) recognized by potential neutralizing antibody in S-protein-ACE2 protein complex (PDB ID: 6M17).

There are three limitations to the SARS-COV-2 proteome microarrays. First, some antibodies may recognize post-translational modifications or conformational (rather than linear) epitopes. To address this issue, we included full-length N, S and E proteins on our microarrays as a comparison. Second, more than one hundred new SARS-COV-2 strains have been identified (https://www.gisaid.org/) since the preparation of our proteome microarray. These new strains could be included in the next version of the SARS-COV-2 proteome microarrays. Third, our SARS-COV-2 proteome microarray is for research use only at this time; it is not approved to diagnose or manage the treatment of patients at this time.

Altogether, our data demonstrate that a peptide-based SARS-COV-2 proteome microarray can map the humoral antibody response to COVID-19. The data could be used to monitor the immune response and identify immunogenic epitopes to develop effective therapeutic treatment of COVID-19. Scientists who wish to acquire these arrays to help fight this COVID-19 pandemic are encouraged to contact us.

## Contributions

X. H., Y. L. X. W. provided the clinical samples. H.W., X. W., X. H., X. Z., T. L., J. D., T. F., S. G., H. D. and X.Y. executed microarray experiments. D. W., X. Z., T. L. and X.Y. executed the bioinformatics and statistical analysis. X.Y., and Y. L. conceived the idea, designed experiments, analyzed the data and wrote the manuscript.

## Acknowledgement

This work was supported by the State Key Laboratory of Proteomics (SKLP-C202001, SKLP-O201703 and SKLP-K201505), the Beijing Municipal Education Commission, National Natural Science Foundation of China (81671618, 81871302, 81673040, 31870823), the National Program on Key Basic Research Project (2018YFA0507503, 2017YFC0906703 and 2018ZX09733003) and the CAMS Initiative for Innovative Medicine (2017-I2M-3-001 and 2017-I2M-B&R-01). We also thank Dr. Brianne Petritis for her critical review and editing of this manuscript.

## Competing interests

None declared.

## Supplementary information

The supplementary information includes the materials and methods, 1 supplementary table and 5 supplementary figures.

## Notes

### Competing Interest Statement

The authors have declared no competing interest.

